# Dynamic regulation of hierarchical heterogeneity in Acute Myeloid Leukemia serves as a tumor immunoevasion mechanism

**DOI:** 10.1101/2020.12.21.414649

**Authors:** Constandina Pospori, William Grey, Sara Gonzalez Anton, Shayin Gibson, Christiana Georgiou, Flora Birch, Georgia Stevens, Thomas Williams, Reema Khorshed, Myriam Haltalli, Maria-Nefeli Skoufou-Papoutsaki, Katherine Sloan, Hector Huerga Encabo, Jack Hopkins, Chrysi Christodoulidou, Dimitris Stampoulis, Francesca Hearn-Yeates, John Gribben, Hans J. Stauss, Ronjon Chakraverty, Dominique Bonnet, Cristina Lo Celso

**Affiliations:** Department of Life Sciences, Faculty of Natural Sciences, Imperial College London, UK; The Francis Crick Institute, London, UK; Centre for Tumour Biology, Barts Cancer Institute, London, UK; Lydia Becker Institute of Immunology and Inflammation, University of Manchester, UK; Wellcome-MRC Cambridge Stem Cell Institute, University of Cambridge; Cancer Research UK Cambridge Institute; Babraham Institute, University of Cambridge; Institute of Ophthalmology, Faculty of Brain Sciences, UCL; Centre for Haematology, Department of Immunology and Inflammation, Faculty of Medicine, Imperial College London; Department of Cell Biology, UCL Institute of Ophthalmology, London, UK; Centre for Haemato-oncology, Barts Cancer Institute, London, UK; Institute of Immunity and Transplantation, University College London; MRC Weatherall Institute of Molecular Medicine, University of Oxford

**Keywords:** AML, tumor heterogeneity, hierarchical heterogeneity, PDL1, LSC, immunotherapies, immunoevasion

## Abstract

Acute Myeloid Leukemia, a hematological malignancy with poor clinical outcome, is composed of hierarchically heterogeneous cells. We examine the contribution of this heterogeneity to disease progression in the context of anti-tumor immune responses and investigate whether these responses regulate the balance between stemness and differentiation in AML. Combining phenotypic analysis with proliferation dynamics and fate-mapping of AML cells in a murine AML model, we demonstrate the presence of a terminally differentiated, chemoresistant population expressing high levels of PDL1. We show that PDL1 upregulation in AML cells, following exposure to IFNγ from activated T cells, is coupled with AML differentiation and the dynamic balance between proliferation, versus differentiation and immunosuppression, facilitates disease progression in the presence of immune responses. This microenvironment-responsive hierarchical heterogeneity in AML may be key in facilitating disease growth at the population level at multiple stages of disease, including following bone marrow transplantation and immunotherapy.

## Introduction

Intratumor heterogeneity is the biggest challenge to the widespread application of precision cancer medicines. Beyond genetic heterogeneity, many cancers also display hierarchical heterogeneity with rare cancer stem cells (CSCs) giving rise to more differentiated progeny (Kreso and Dick, 2014). Clearing quiescent, chemoresistant CSCs is critical for achieving cure yet remains difficult to achieve using current approaches. At the same time, because the differentiated progeny of CSCs is generally considered unable to propagate disease, its contribution to disease evolution remains understudied. It is unknown whether the generation of differentiated malignant cells may be regulated by dynamic changes in the tumor microenvironment (TME) or whether these differentiated cells may have important functional properties, for example shaping the TME to make it more supportive of CSC survival.

Acute Myeloid Leukemia (AML) has been a paradigm for the CSC hypothesis, following the phenotypic identification of leukemic populations (leukemia stem cells, LSCs) able to recapitulate the human disease in mouse xenografts (Bonnet and Dick, 1997). AML typically presents as perturbed hematopoiesis in which excessive proliferation and a block in differentiation drive the accumulation of myeloid cells at a range of differentiation stages (Dohner et al., 2015). Therefore, AML provides an ideal platform to study the relevance of hierarchical heterogeneity to cancer progression.

In AML patients, conventional chemotherapy still leads to overall poor survival rates because of disease relapse driven by chemoresistant cells. Allogeneic hematopoietic stem cell transplantation leads to cure in a portion of patients due to the graft-vs-leukemia response (Dickinson et al., 2017). Despite efforts to manipulate donor HSCs to improve transplantation outcomes (Fares et al., 2014; Goncalves et al., 2016; Grey et al., 2020; Holmfeldt et al., 2016), accumulating data show that post-transplant relapse correlates with T cell exhaustion or immune-edited variants lacking major histocompatibility antigen expression (Christopher et al., 2018; Noviello et al., 2019; Toffalori et al., 2019). Immunotherapies have been very promising for malignancies such as melanoma (Leonardi et al., 2020; Luke et al., 2017) and B-cell Acute Lymphoblastic Leukemia (Ghorashian et al., 2019; Maude et al., 2018; Park et al., 2018), but have so far produced disappointing results against AML. Although anti-tumor immune responses have been reported in a variety of AML murine models and in AML patients (Hayashi et al., 2019; Rezvani et al., 2005; Zhang et al., 2009; Zhou et al., 2011), paucity of AML-specific antigens, limited *in vivo* persistence of adoptively transferred T cells, and inability to precisely identify patients likely to benefit have limited immunotherapeutic approaches (Witkowski et al., 2019). Thus, a deeper understanding of the interactions between AML cells and immune cells is needed to improve the efficacy of immunotherapies.

The hierarchically heterogeneous malignant myeloid cells composing AML are likely to maintain diverse immune-modulating functions. Recent work combining single cell transcriptomic and genomic analyses of bone marrow (BM) samples from AML patients and healthy donors has demonstrated the presence of differentiated, monocyte-like leukemic cells with the capacity to suppress T cells *in vitro* (van Galen et al., 2019). While high AML LSC frequency correlates with worse patient prognosis (Ng et al., 2016; Thorsson et al., 2018), it is unclear if AML hierarchical heterogeneity has a role in facilitating disease progression and relapse. Furthermore, although extensive work has demonstrated that inflammation rewires healthy hematopoiesis to orchestrate a shift in the balance between stemness and differentiation (Baldridge et al., 2010; Batsivari et al., 2020; King and Goodell, 2011; Kovtonyuk et al., 2016), it remains unclear whether malignant hematopoiesis would be similarly affected.

Here, we dissect the reciprocal links between anti-tumor immune responses and hierarchical heterogeneity in AML. We identify T cell responses against AML, and as induction of PDL1 upregulation on tumor cells by T cell-derived IFNγ (Garcia-Diaz et al., 2017; Spranger et al., 2013) is a key cancer immunoevasion mechanism in a wide variety of cancers (Sun et al., 2018), we examine PDL1 expression on AML cells. Although it has previously been shown that IFNγ can induce PDL1 expression in AML patient samples (Berthon et al., 2010; Kronig et al., 2014), the clinical importance of this checkpoint molecule in AML remains controversial (Antohe et al., 2020; Berthon et al., 2010; Chen et al., 2008; Ghosh et al., 2020; Salih et al., 2006; Tamura et al., 2005). We demonstrate here that PDL1 expression levels on AML cells are dynamic during disease progression and that the highest expression is seen among terminally differentiated AML cells. We show that PDL1 upregulation in response to anti-tumor T cell responses is associated with a shift in AML cell fate towards increased differentiation and reduced stemness. Despite the reduced intrinsic leukemia propagating capacity, these heterogeneous AML cell populations have a growth advantage in the context of active antitumor immune responses.

## Results

### Dynamic T cell responses in the BM microenvironment as AML progresses

To investigate the effect of anti-tumor immune responses on AML lineage hierarchy, we used the well-established and clinically relevant MLL-AF9 transduction-driven model of murine AML, in which the c-Kit^+ve^ AML subpopulation is enriched for LSCs (Krivtsov et al., 2006; Somervaille and Cleary, 2006). AML cells harvested from fully infiltrated primary mice (Duarte et al., 2018) were transplanted into secondary fully immunocompetent syngeneic C57Bl/6 mice or immunodeficient mice. In some experiments, leukemic mice or harvested leukemic cells *ex vivo* were treated with chemotherapy or leukemia-specific T cells (Figure 1A). As expected, disease was first detected in BM and subsequently the spleen (Duarte et al., 2018) and blood of injected mice (Figure 1B and C).

**Figure 1.**
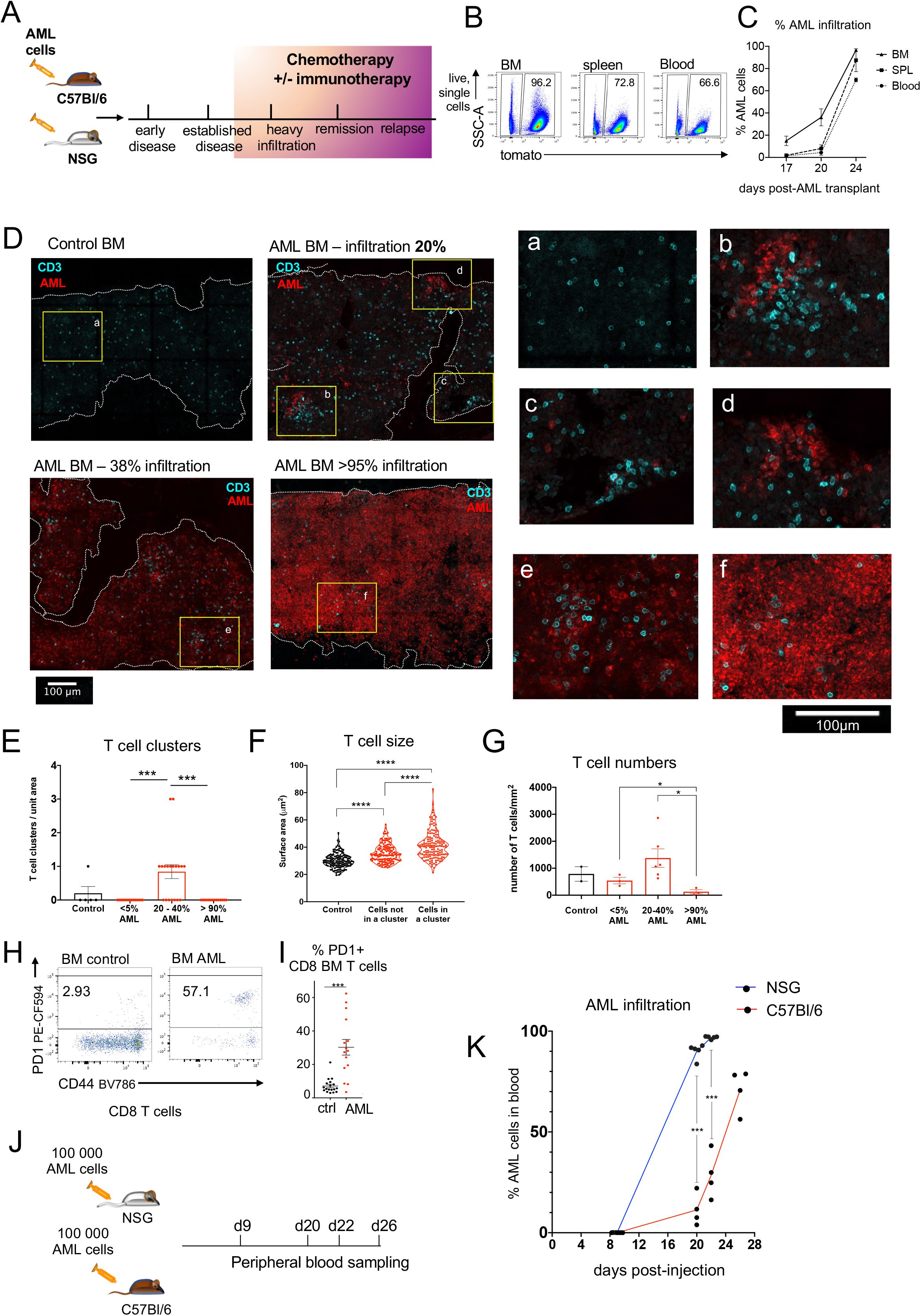
Dynamic T cell changes take place in the BM microenvironment as AML progresses. (A) Schematic diagram of the experimental setup. Up to 100 000 AML cells were injected in C57Bl/6 mice or immunodeficient mice, as specified in individual experiments. Disease progression and AML cells phenotype were monitored longitudinally in the blood of transplanted mice or in the BM, spleen and blood of mice euthanised at specific timepoints. Transplanted mice received chemotherapy in some experiments, whereas BM and spleen cells were harvested and treated with chemotherapy or leukemia-specific cytotoxic T cells *in vitro* in other experiments. (B) 100 000 primary, tomato FP+ cells were injected IV, in C57Bl/6 recipients. BM, spleen and blood were harvested 24 days after-injection and leukemic cells were detected in each tissue by FACS. (C) Graph summarising the percentage of AML infiltration in BM, spleen and blood of leukemic mice, at d17, 20, and d24 from same set up as in (B). n=3 mice per time point. Lines connect the mean AML infiltration in each tissue, at each time point. Error bars: mean ± SEM. (D) Representative maximum intensity projections of tilescan images of 20 μm thick sections of tibias or sternums from control and leukemic C57Bl/6 mice with progressive disease infiltration, stained with anti-CD3 antibody to identify T cells. Leukemic cells were identified through the expression of tomato FP or YFP. (a-f) inserts show higher resolution views of the areas indicated in the tilescans. (E) Number of T cell clusters/0.25mm^2^ area observed in control mice and in leukemic mice at different levels of AML infiltration in the BM. Each dot represents an individual area sampled. Mann-Whitney test was used for significance, error bars: mean ± SEM. (F) Individual T cells’ size in control BM and in BM with up to 40% overall leukemia infiltration. T cells from leukemic BM are categorised according to their localisation within clusters or not. Each dot represents an individual T cell, and 150 T cells per category (healthy control BM tilescans, T cell in clusters and T cells not in a cluster in 20-40% AML infiltration tilescans) were sampled. T cells in clusters/not in clusters were sampled in paired areas from 3 leukemic mice. Mann-Whitney test used for significance. (G) Graph summarising the T cell numbers/mm^2^ in 20 μm BM sections. Each dot represents data from an individual animal. Students t-test used for significance, error bars: mean ± SEM. D-G: Data pooled from 3 independent experiments. Controls n=2 mice, infiltration<5% n=3 mice, infiltration 20-40% n= 6 mice, infiltration> 95% n= 3 mice. (H) Representative FACS plots showing PD1 and CD44 expression on BM CD8+ T cells in control and in fully infiltrated C57Bl/6 mice. (I) Graph showing the percentage of BM CD8 T cells expressing PD1 in control and in fully infiltrated C57Bl/6 mice. H and I: n= 18 control and n= 15 leukemic C57Bl/6 mice, pooled from 3 independent experiments. Student’s t-test used for significance, error bars: mean ± SEM. (J) Schematic diagram of experimental setup in which 100 000 AML cells were injected into age-matched C57Bl/6 and NSG recipients. Blood was sampled at 9, 20, 22 and 26 days after-injection to monitor disease infiltration. All NSG mice had to be sacrificed by d22 due to disease progression. (K) AML infiltration in peripheral blood in NSG and in C57Bl/6 mice described in (J). Each dot represents an individual mouse, lines connect mean infiltration at each time point, in each group. Multiple t-tests used for significance with Holm-Sidac correction. NSG mice n=5, C57Bl/6 mice n=4.

Anti-CD3 antibody immunostaining of BM sections from healthy and AML burdened C57Bl/6 mice revealed dynamic changes in BM T cell distribution as AML cells grew. In contrast to healthy BM where T cells were evenly distributed (Figure 1D-*a*), we observed clustering of T cells in leukemic mice, either directly adjacent to initial foci of AML cells (Figure 1D-*b* and Figure 1E, Figure S1) or in positions distant from AML cells (Figure 1D-*c* and Figure 1E, Figure S1). The size of T cells within clusters was larger than that of T cells not in clusters (Figure 1F), suggesting that T cells were responding to AML cells locally and may have even cleared them from some areas. However, T cell clusters were often absent from areas containing AML foci (Figure 1D-*d*). As disease progressed, T cells were largely excluded from BM areas where large AML patches had established, and the identifiable T cell clusters appeared to be less compact (Figure 1D-e). Eventually, in fully infiltrated mice, T cell clusters were no longer present in BM (Figure 1D-f and 1E) and the overall number of BM T cells was reduced compared to control animals (Figure 1D-a and f and 1G).

FACS analysis of BM from control and fully infiltrated mice showed a high percentage of CD8 T cells expressing both PD1 and CD44 (Figure 1H and I), a phenotype associated with tumorinfiltrating lymphocytes (Gros et al., 2014), suggesting an anti-tumor T cell response occurred in leukemic mice, albeit ineffective. Longitudinal sampling of the blood of C57Bl/6 and NSG mice injected with 100 000 AML cells (Figure 1J) showed that the disease grew faster in the latter (Figure 1K), implying that anti-tumor immune responses initially controlled AML growth.

### Dynamic changes in AML phenotype during disease progression and following treatments

The evidence of T cell activation in response to tumor raised the possibility that the ensuing inflammation could influence hierarchical heterogeneity in AML. We examined PDL1 and c-Kit expression levels among leukemic cells, as surrogate markers of their immunosuppressive and leukemia propagating capacity (LPC) respectively. Typically, very few cells from fully infiltrated primary mice, used for the secondary transplantation, were PDL1hi (Figure 2 A). PDL1 expression was highest among c-Kit^-ve^ cells (Figure 2A and 2B), known to consist of more differentiated AML cells (Saadatpour et al., 2014; Somervaille and Cleary, 2006). Intriguingly, longitudinal analysis of leukemic mice peripheral blood showed that the majority of AML cells at early-intermediate disease (<15% AML infiltration in blood) expressed high levels of PDL1, with the ones expressing the highest levels of PDL1 being c-Kit^-ve^. With disease progression, PDL1 expression levels diminished among most AML cells, and PDL1^hi^c-Kit^-ve^ and PDL1^lo^c-kit^+ve^ populations were identifiable (Figure 2B and 2C). The PDL1^hi^ c-Kit^-ve^ population was always more prominent among AML cells in blood compared to those present in spleen and BM (Figure 2D).

**Figure 2.**
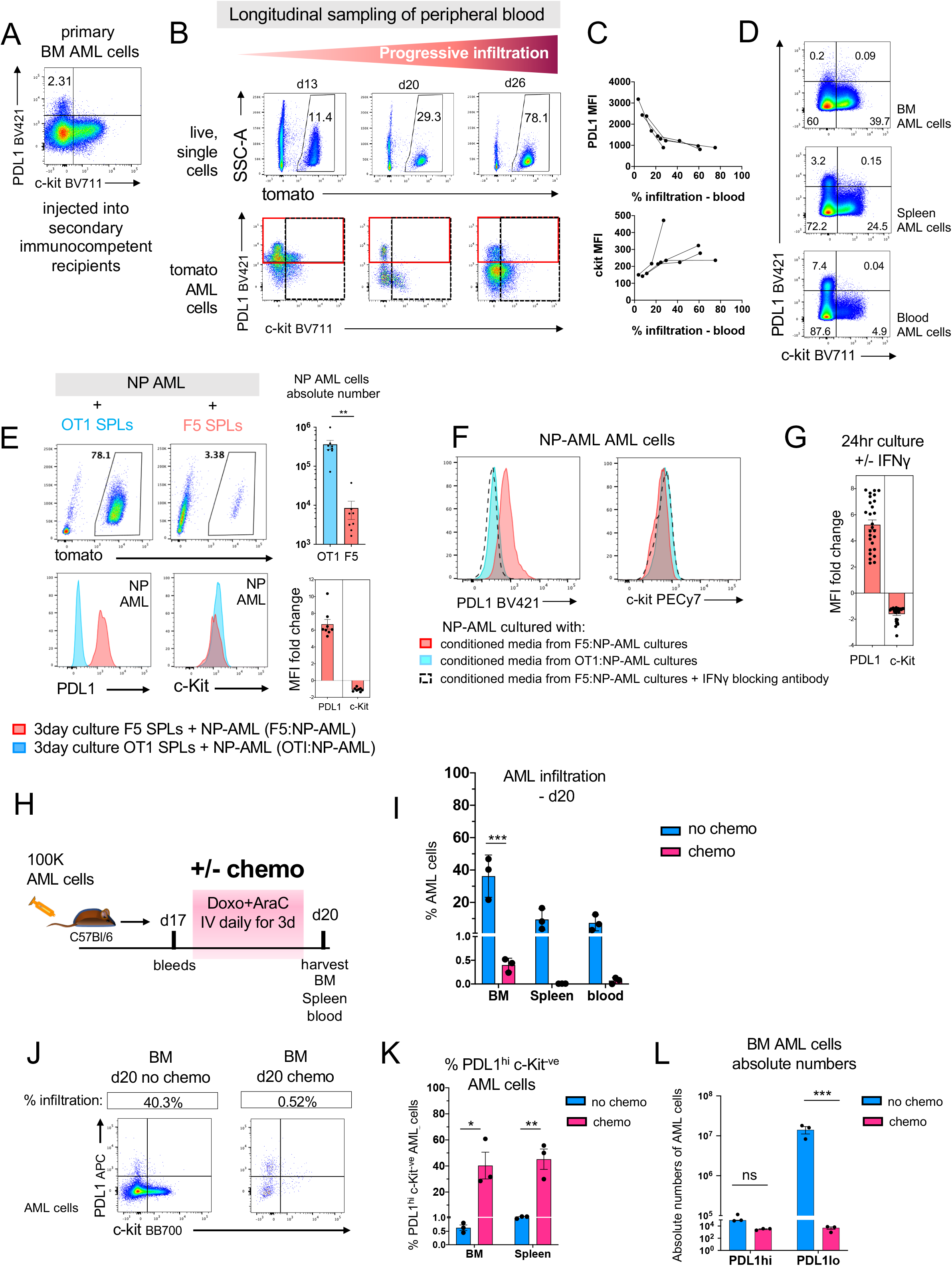
Dynamic changes in AML phenotype during disease progression and following treatments. (A) PDL1 and c-Kit expression on BM AML cells from primary C57BL/6 leukemic mice prior to injecting them into secondary C57Bl/6 recipients. (B) C57Bl/6 mice were injected with 100 000 AML cells and blood was sampled at d13, d20 and d26 and stained for PDL1 and c-Kit expression. Red squares highlight PDL1^hi^ cells. Representative of 3 independent experiments. (C) PDL1 and c-Kit Mean Fluorescent Intensity in relation to % AML infiltration in the blood, from experiment in (B). n= 4 mice. Each line links data from an individual mouse. (D) Representative FACS plots showing PDL1 and c-Kit expression on AML cells in BM, spleen and blood. (AML infiltration: 87.5% in BM, 62.6% in spleen and 84.6% in blood). Data representative of 3 independent experiments. (E) FACS analysis of NP-AML cells co-cultured with F5 or OTI TCRtg splenocytes (1:2 ratio) for 3 days. Top row shows SSC-A vs tomato FP expression among all live cells in the cultures, with NP-AML cells identified as being tomato FP^+ve^. Bottom row shows overlay histograms of PDL1 (left) and c-Kit expression (right) on NP-AML cells cultured in the presence of F5 (red) or OTI TCRtg splenocytes (blue). Top graph summarises absolute number of live NP-AML cells following 3 days in culture with OTI or F5 splenocytes. Bottom graphs summarise fold change in PDL1 and c-Kit MFIs, for each NP-AML sample in F5 co-cultures, compared to OTI co-cultures. n= 8 OT1 and F5 independent co-cultures. Error bars: mean ± SEM. Paired t test used for significance. (F) PDL1 and c-Kit expression on secondary NP-AML cells cultured for 3 days in conditioned media from F5:NP-AML and OTI:NP-AML co-cultures in (E), and in F5:NP-AML co-culture conditioned medium supplemented with IFNγ blocking antibody. Data shown are representative of technical triplicates. (G) Fold differences in PDL1 and c-Kit MFI in BM and spleen AML cells cultured for 24hrs in the presence or absence of 10ng/ml IFNγ. n= 27 independent cultures (data pooled from 3 independent experiments) Error bars: mean ± SEM. (H) Schematic diagram of experimental setup. 100 000 AML cells were injected in C57Bl/6 recipients and blood was monitored for AML infiltration. At d17, when less than 1% AML was detected in the blood, mice were either left untreated or received a combination of cytarabine and doxorubicin intravenously for 3 days. Mice were sacrificed on d20 and BM, spleen and blood were harvested and PDL1 and c-Kit expression on AML cells was analysed by FACS. Untreated leukemic mice, n=3. chemotherapy treated mice, n=3. (I) AML infiltration in BM, spleen and blood of mice 20 days post-AML injection, untreated or treated with cytarabine and doxorubicin (described in H). Each dot represents an individual mouse, error bars: mean ± SEM, multiple t tests used for significance. (J) Representative FACS plots of PDL1 and c-Kit expression on AML cells from untreated and chemotherapy-treated leukemic mice (described in H). (K) Percentage of PDL1^hi^ c-Kit^-ve^ AML cells among all AML cells in BM and in spleen of untreated and chemotherapy-treated mice (described in H). Each dot represents data from individual mice, error bars: mean ± SEM, multiple t-tests with Holm-Sidak correction, performed for statistical significance. (L) Absolute numbers of PDL1^hi^ and PDL1^lo^ AML cells in the BM of untreated and chemotherapy-treated mice (described in H). Each dot represents data from individual mice, error bars: mean ± SEM, multiple t-tests with Holm-Sidak correction, performed for statistical significance.

To test if an anti-tumor T cell response could induce the phenotype dynamics observed among circulatory leukemic cells, AML cells expressing the influenza A nucleoprotein (NP) model-antigen were generated and grown in Rag1 KO and NSG immunodeficient mice (Figure S2). NP-AML cells were subsequently harvested from BM and spleen of these animals and were challenged *in vitro* with F5 or OTI TCR transgenic (tg) splenocytes, containing NP-specific or control OVA-specific CD8 T cells, respectively. In these co-cultures, NP-AML cells grew unaffected by OTI T cells but were efficiently killed by F5 T cells (Figure 2E top row). The few NP-AML cells surviving the F5 T cell response expressed higher levels of PDL1 and lower levels of c-Kit compared to the NP-AML cells co-cultured with OTI T cells (Figure 2E bottom row). This phenotype was similar to that observed *in vivo* in blood AML cells, at early-intermediate disease (Figure 2B). Culturing AML cells with conditioned media from the F5:NP-

AML co-cultures, in the presence or absence of an IFNγ blocking antibody showed that the shift towards a PD-L1^hi^ c-Kit^lo^ phenotype was IFNγ-dependent (Figure 2F). Culturing AML cells with 10ng/ml IFNγ, for 24hrs was sufficient to recapitulate PDL1 upregulation and c-Kit downregulation (Figure 2G).

Next, PDL1 and c-Kit expression levels were assessed on chemoresistant AML cells. Intravenous (IV) injection of a combination of cytarabine plus doxorubicin, daily for 3 days, initiated at an early disease infiltration (average infiltration = 0.3% in peripheral blood, d17 post-AML injection) (Figure 2H) cleared the leukemia from blood, while only 0.005% and 0.4% AML infiltration persisted in the spleen and the BM respectively. Untreated leukemic mice progressed to higher infiltration in all 3 organs studied (Figure 2I). The PDL1^hi^ population was highly enriched among the BM AML cells in treated animals compared to those in untreated animals, where PDL1^hi^ AML cells remained rare (Figure 2J and 2K). The loss of PDL1^lo^ AML cells in absolute numbers was significantly higher than that of PDL1^hi^ AML cells, with the latter appearing to be essentially chemoresistant (Figure 2L). Longitudinal sampling of peripheral blood in leukemic mice treated with cytarabine alone, at a later stage of disease progression, also showed the decrease in disease load to be coupled with an enrichment in PDL1^hi^ leukemic cells (Figure S3). Taken together, these data indicate that the phenotype of AML cells is dynamic following both immune responses and chemotherapy and raise questions about the proliferative, LPC and immunomodulatory function of the PDL1^hi^ AML cells, and about the lineage relationship between PDL1^hi^ and PDL1^lo^ AML cells.

### PDL1^hi^ AML cells are non-proliferative

To investigate the proliferation profile of AML cell subpopulations *in vivo*, we performed EdU/BrdU dual pulse chase experiments (Akinduro et al., 2018) which allowed the identification of leukemic cells at different cell cycle stages in a 2-hr window between the EdU pulse and the BrdU pulse. The four cell subpopulations are: EdU^-ve^-BrdU^-ve^ (DN) AML cells in G0 or G1; EdU^+ve^-BrdU^+ve^ (DP) AML cells in S-phase throughout the dual-pulse chase (late S-phase); EdU^-ve^ BrdU^+ve^ (BrdU SP) AML cells that entered S-phase in the 2-hr window following EdU administration (early S-phase); and EdU^+ve^ BrdU^-ve^ (EdU SP) AML cells containing cells that had completed S phase prior to the BrdU pulse. To distinguish between AML cells that transitioned to G2-M from those that completed mitosis and divided in the 2hr window within the EdU SP population, EdU dilution vs FSC-A was assessed; cells in G2-M had higher EdU MFI and higher FSC-A, and cells that had divided, had lower EdU MFI and lower FSC-A (Figure 3B, 3C and 3D).

**Figure 3.**
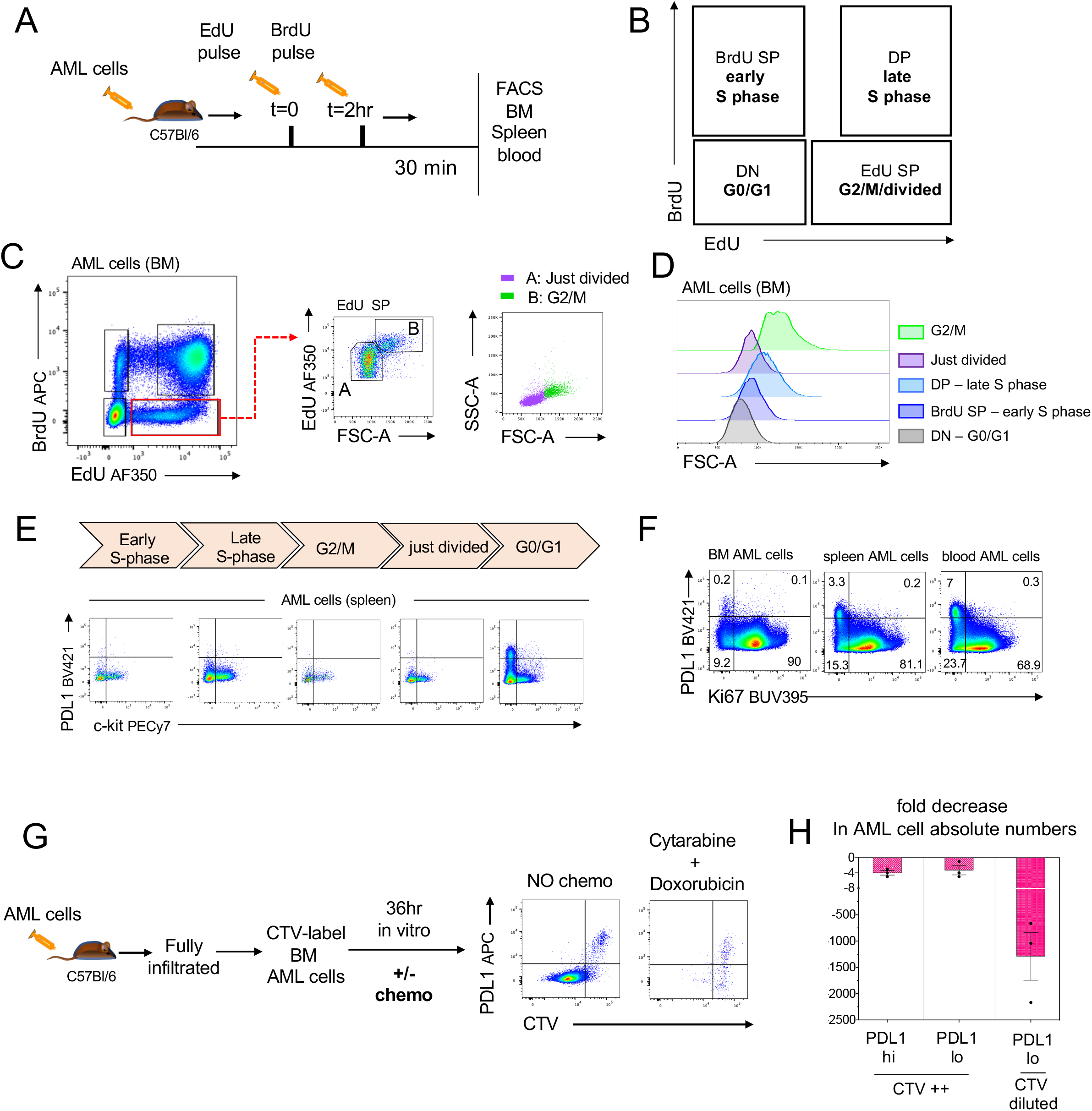
Assessing proliferation dynamics of phenotypically heterogeneous AML cells. (A) Schematic diagram of experimental set up. Leukemic mice were injected with a single dose of EdU followed by a single dose of BrdU 2hrs later. Mice were euthanised 30min after the BrdU pulse and BM, spleen and blood were harvested and FACS analysed for nucleoside analogue incorporation as well as for PDL1 and c-Kit expression. (B) Schematic diagram describing the FACS gates corresponding to cell cycle stages in the 2hr pulse-chase window. (C) Representative FACS plots of EdU/BrdU incorporation in BM AML cells (left panel) and EdU vs FSC-A among the EdU SP AML population (middle panel). The latter identifies 2 populations (A: lower EdU MFI, lower FSC-A just divided cells; B: higher EdU MFI, higher FSC-A G2-M cells). A and B populations overlaid in FSC-A-SSC-A FACS plots (right panel). (D) FSC-A histogram overlays of BM AML cells at each cell cycle stage, as identified in (C). (E) Representative FACS plots showing PDL1 and c-Kit expression on spleen AML cells at each cell cycle stage as specified. C-E data from 3 independent experiments. n= 9 leukemic mice. (F) Representative FACS plots showing PDL1 and Ki67 staining on BM, spleen and blood AML cells in leukemic mice. 2 independent experiments, n=10 mice. (G) Schematic diagram of experimental set up. BM AML cells harvested from leukemic mice were Cell Trace Violet (CTV)-labelled and cultured in the presence or absence of chemotherapy (cytarabine and doxorubicin). 36hr later, the cells were FACS analyzed for PDL1 expression and CTV dilution and representative FACS plots are shown. (H) Graph summarising the fold-decrease in the absolute numbers of PDL1^hi^-CTV bright, PDL1^lo^-CTV bright and PDL1^lo^-CTV diluted cells following 36hr of culture in the presence of chemotherapy. n=3 leukemic mice, error bars: mean ± SEM. One-way ANOVA indicated statistical significance (p=0.038). Data representative of 2 independent experiments.

AML cells at each of these cell cycle stages were analyzed for expression of PDL1 and c-Kit (Figure 3E). Notably, PDL1^hi^ c-Kit^-ve^ AML cells were only present among the DN population in BM, spleen and blood (Figure 3E and not shown). As the EdU SP FSC-A^lo^, ‘just divided’, population did not contain PDL1^hi^ c-Kit^-ve^ AML cells (Figure 3E, fourth panel), it appears that the increase in PDL1 expression on leukemic cells is not an immediate result of proliferation. In separate experiments, PDL1^hi^ AML cells were shown to be Ki67^-ve^ in BM, spleen and blood (Figure 3F). These data indicated that PDL1^hi^ AML cells were in G0.

The non-proliferative state of PDL1^hi^ AML cells was consistent with chemoresistant leukemic cells being enriched for this subset *in vivo* (Figure 2J, 2K, 2L). Analyzing ex-vivo Cell Trace Violet (CTV)-labelled AML cells kept in culture for 36hr, in the presence or absence of chemotherapy, showed that the PDL1^hi^ AML cells did not dilute CTV, confirming that they were non-proliferative under both conditions (Figure 3G). A small proportion of PDL1^lo^ AML cells also appeared to be non-proliferative (CTV^hi^). PDL1^hi^ and PDL1^lo^ non-proliferative AML cells were similarly and minimally susceptible to chemotherapy, while the opposite was the case for proliferative AML cells (Figure 3H).

### PDL1^hi^ AML cells are terminally differentiated progeny of PDL1^lo^ AML cells

To determine if PDL1^hi^ AML cells are the terminally differentiated progeny of PDL1^lo^ AML cells, we performed a lineage tracing experiment using an EdU pulse-chase. Leukemic mice at early infiltration (d17, average blood infiltration=0.8%) were injected with a single dose of EdU and peripheral blood was sampled 20hr, 48hr, and 6 days later to trace the fate of AML cells that had incorporated EdU (Figure 4A). Consistent with the non-proliferative state of PDL1^hi^ AML cells, EdU was only detectable in PDL1^lo^ AML cells at 20hr post-EdU pulse (Figure 4B). At the 48hr timepoint, the MFI of EdU^+ve^ AML cells was reduced compared to that at the 20hr timepoint (Figure 4B and 4C) indicating that the initially labelled cells had proliferated. At this same time point, PDL1^hi^ EdU^+ve^ AML cells started appearing (Figure 4B and 4E). In agreement with our earlier findings of a transient increase in PDL1 expression among circulating AML cells in early/intermediate disease (Figure 1B), the PDL1 MFI on AML cells here was the highest at the 48hr time point (Figure S4A) when the percentage AML infiltration in the blood ranged between 1 and 2.7%. By 6 days post-EdU pulse (d23 post-AML injection), all PDL1lo AML cells that initially incorporated EdU had lost their EdU label (Figure 4B and 4D) while a population of PDL1^hi^-EdU^dim^ AML cells was clearly present, confirming the lineage relationship between PDL1lo (progenitor) and PDL1hi (progeny) AML cells (Figure 4B and 4E). Consistent with this, FACS sorted, PDL1^lo^ c-Kit^+ve^ progenitor-like AML cells transplanted into C57Bl/6 mice generated all three main AML subsets PDL1^hi^ c-Kit^-ve^, PDL1^lo^ c-Kit^-ve^, and PDL1^lo^ c-Kit^+ve^ (Figure S4B and S4C). Importantly, in this serial transplant setting, we observed again the transient enrichment in PDL1^hi^ leukemic cells among circulatory AML cells in early/intermediate disease (Figure S4D).

**Figure 4.**
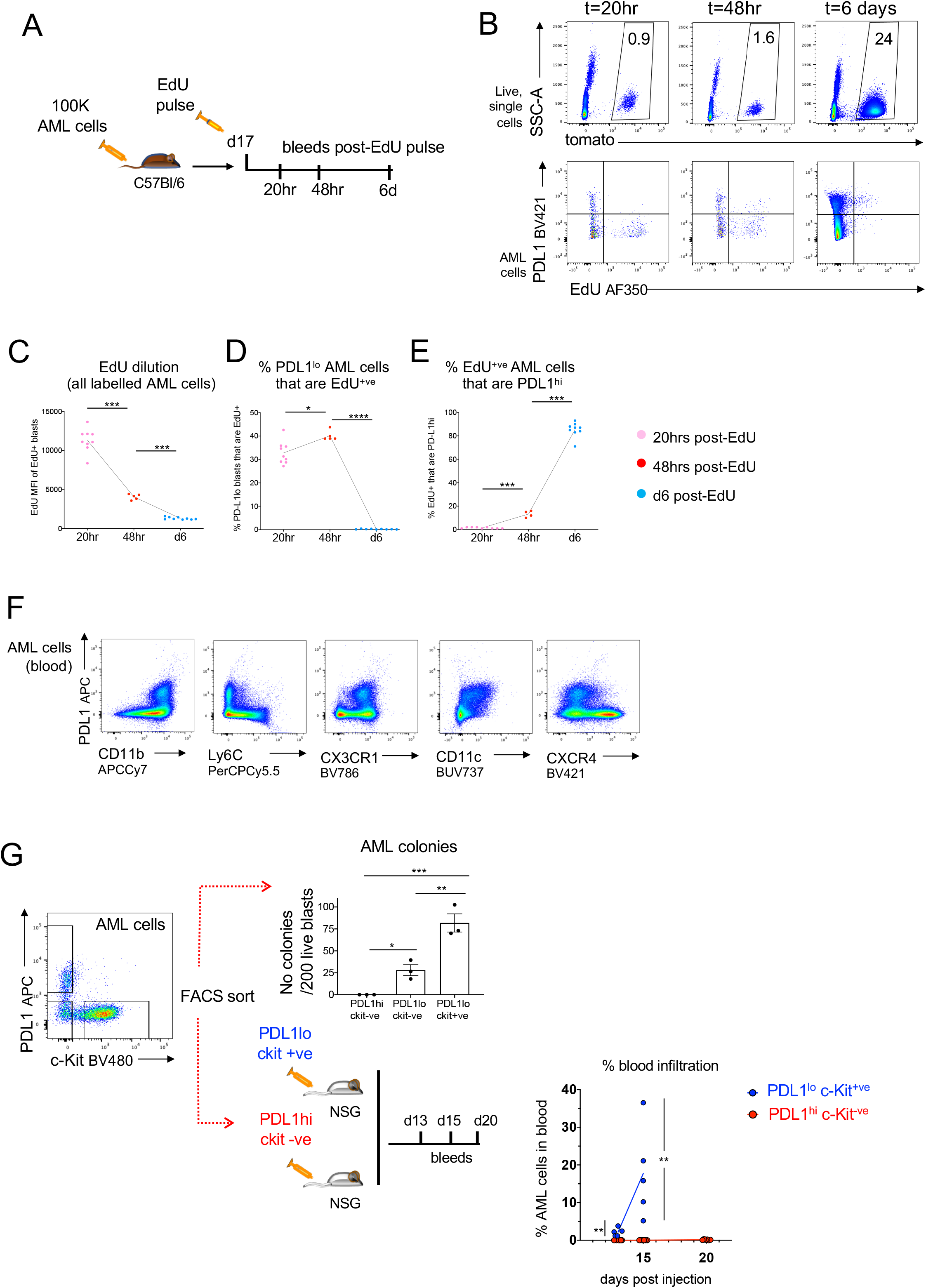
Lineage relationship between PDL1^lo^ and PDL1^hi^ AML cells. (A) Schematic diagram of experimental setup. Leukemic mice were intravenously injected with a single pulse of EdU 17 days post-AML injection when blood AML infiltration was less than 1%. EdU labelling of AML cells and EdU dilution in the blood was monitored at 20hrs, 48hrs and 6 days post-EdU pulse. (B) Representative FACS plots of blood samples from a leukemic mouse injected with a single dose of EdU (as described in A) at the 20hr, 48hr and 6-day timepoints. Top row shows AML infiltration. Bottom row shows EdU labelling in relation to PDL1 expression in AML cells. n=9 C57Bl/6 leukemic mice. (C) EdU MFI among all EdU^+ve^ AML cells at 20hr, 48hr and d6 post-EdU pulse. Each dot represents data from individual mice. Paired t-tests used for statistical significance. (D) % of PDL1^lo^ AML cells that are EdU^+ve^ at 20hr, 48hr and d6 post-EdU pulse. Each dot represents data from individual mice. Paired t-tests used for statistical significance. (E) % of EdU^+ve^ AML cells that are PDL1^hi^ at 20hr, 48hr and d6 post-EdU pulse. Each dot represents data from individual mice. Paired t-tests used for statistical significance. (F) Blood harvested from fully infiltrated leukemic mice was stained for markers expressed on differentiated myeloid cells, including CD11b, Ly6C, CX3CR1, CD11c and CXCR4. n=4 leukemic mice. Data representative of 3 independent experiments. (G) Blood was harvested from 3 fully infiltrated leukemic C57Bl/6 mice and AML cells were sorted based on PDL1 and c-Kit expression and plated in colony forming assays. Each dot represents the average number of colonies per 200 sorted AML cells from each leukemic mouse. Paired t-tests were used for statistical significance, error bars: mean ± SEM. Separately, blood was pooled and 5000 sorted PDL1^hi^c-Kit^-ve^ or PDL1^lo^c-Kit^+ve^ AML cells were injected intravenously in NSG mice to assess their leukemia propagating capacity. Blood was sampled at d13, d15 and d20 following AML cell injection to monitor disease progression. The mice receiving PDL1^lo^ c-Kit^+ve^ AML cells were fully infiltrated and had to be sacrificed at the day15 timepoint. n=5 NSG mice received PDL1^lo^c-Kit^+ve^ AML cells, n=6 NSG mice received PDL1^hi^c-Kit^-ve^ AML cells. Each dot represents data from individual mice, lines connect mean infiltration at each time point, in each group. T-tests with Holm-Sida correction, at each time point used for statistical significance

Next, we went on to characterize the phenotype of PDL1^hi^ and PDL1^lo^ AML cells more extensively (Figure 4F). PDL1^hi^ AML cells expressed high levels of CD11b and, interestingly, were also Ly6C^-ve^ and CX3CR1^+ve^. This pattern of surface marker expression indicated an overlapping phenotype of PDL1^hi^ AML cells with that of terminally differentiated, patrolling monocytes (Bianchini et al., 2019). Of note, among PDL1^lo^ AML cells, a range of expression levels is observed for CD11b, Ly6C and CX3CR1 suggesting that differentiated populations are not exclusive to the PDL1^hi^ AML cell compartment. PDL1^hi^ cells expressed low levels of CD11c, and PDL1^lo^ cells were largely negative. Finally, PDL1^hi^ AML cells expressed low levels of CXCR4 compared to PDL1^lo^ cells, which had both CXCR4 high and low subpopulations. Low levels of CXCR4 could contribute to PDL1^hi^ AML cells not being efficiently retained in the BM and spleen and being found most abundant in peripheral blood (Figure 2D).

*In vitro* colony-forming capacity (CFU-C) assays, performed with AML cells sort-purified based on PDL1 and c-Kit expression, proved that PDL1^hi^c-Kit^-ve^ cells have no clonogenic capacity (Figure 4G, top). Moreover, injecting equal numbers of PDL1^hi^ c-Kit^-ve^ and PDL1^lo^ c-Kit^+ve^ AML cells in NSG mice, confirmed that the former were unable to propagate disease *in vivo* (Figure 4G, bottom). These data were consistent with the PDL1^hi^c-Kit^-ve^ AML cell phenotype distinctly marking a non-proliferative, terminally differentiated population.

### IFNγ biases the AML differentiation trajectory to generate a PDL1^hi^ differentiated population

Knowing that IFNγ leads to higher expression of PDL1 on AML cells (Figure 2E-G), and that PDL1^hi^ cells are differentiated cells with no LPC (Figure 4), we next asked whether T cell-derived IFNγ would affect the functional properties of AML cells. To do so, we exposed unfractionated AML cells to conditioned media from F5:NP-AML or OTI:NP-AML co-cultures, in the presence or absence of an IFNγ blocking antibody and assessed their clonogenic capacity in CFU-C assays. AML cells exposed to F5:NP-AML medium generated fewer colonies, but this was rescued by blocking IFNγ (Figure 5A). Similarly, plating unfractionated AML cells in semisolid methylcellulose media supplemented with 10ng/ml IFNγ significantly decreased their CFU-C potential (Figure 5B). Interestingly, the majority of colonies seen in the IFNγ-supplemented cultures had fewer cells and appeared flatter and less compact than the AML colonies growing in methylcellulose media alone (Figure 5C). The appearance of the colonies was consistent with increased differentiation and loss of leukemia propagating potential following exposure to IFNγ. Treating AML cells with IFNγ prior to plating them on methylcellulose media alone resulted in a similar decrease in CFU-C capacity (Figure 5D).

**Figure 5.**
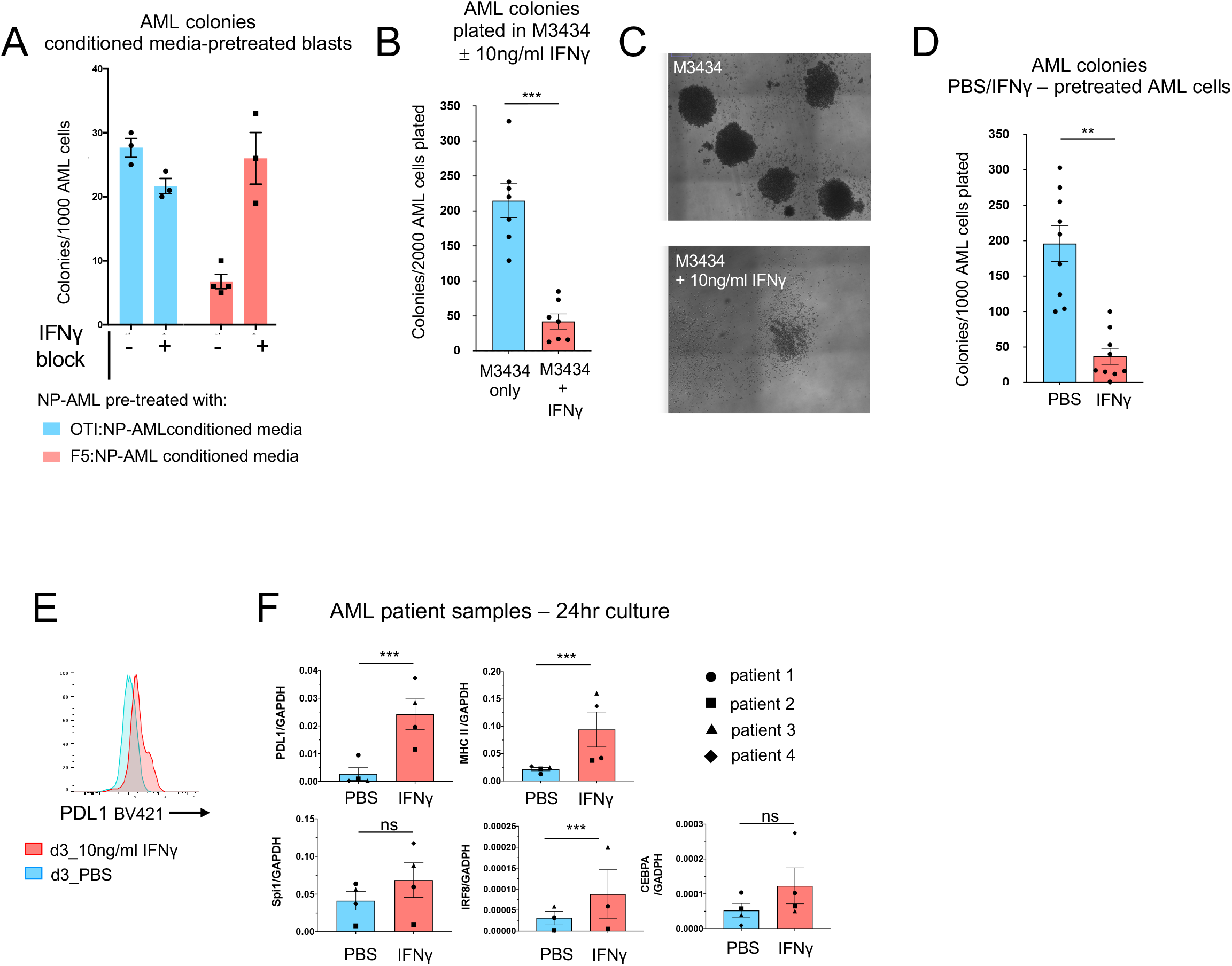
IFNγ-induced PDL1 upregulation on AML cells correlates with a decrease in leukemia propagating capacity. (A) AML cells were cultured for 3 days in conditioned media (1:1 volume) from F5:NP-AML cocultures, or OTI:NP-AML cocultures, in the presence or absence of IFNγ blocking antibody and subsequently plated in colony forming assays. Each dot represents number of colonies generated from technical replicates. error bars: mean ± SEM. (B) Graph showing number of colonies generated from plating 2000 BM or spleen AML cells in colony forming assays, in the presence of 10ng/ml IFNγ or PBS control. Each dot represents colonies grown from AML cells harvested from individual leukemic mice. Paired t-test performed for statistical significance, error bars: mean ± SEM. (C) Representative images of AML cells colonies growing in M3434 medium alone or supplemented with 10ng/ml IFNγ. (D) Number of colonies generated from plating 1000 BM or spleen AML cells pre-treated with PBS or 10ng/ml IFNγ for 24hrs in colony forming assays. Each dot represents colonies grown from AML cells harvested from individual leukemic mice. Paired t-test performed for statistical significance. Error bars: mean ± SEM (E) Representative PDL1 expression in one of 4 AML patient samples following a 3-day culture in the presence of 10ng/ml hIFNγ or PBS. (F) qPCR analysis for *PDL1, MHC II, Spi1, IRF8* and *CEBPA* expression was performed in patient samples following a 24hr culture, in the presence of 10ng/ml hIFNγ or PBS. Gene expression relative to *GAPDH* housekeeping gene shown. Paired t-tests performed for statistical significance. Error bars: mean ± SEM.

PDL1 upregulation following *in vitro* treatment with IFNγ has previously been reported in AML patient samples (Berthon et al., 2010; Kronig et al., 2014). To assess if human AML samples, similarly to murine MLL-AF9-driven AML, are primed to differentiate upon IFNγ exposure, we cultured 4 primary AML samples (table S1) in the presence of 10ng/ml IFNγ or PBS. FACS analysis after 3 days of *in vitro* culture confirmed IFNγ-induced PDL1 upregulation (Figure 5E). qPCR analysis after a 24hr culture in the same conditions, showed an increase in *PDL1* mRNA and, consistent with recent studies (Christopher et al., 2018), increased *MHC II* gene expression in response to IFNγ (Figure 5F). PDL1 upregulation was coupled with a trend for increased expressions of *Spi1* (encoding the Pu.1 transcription factor) and *CEBPA* and a significant upregulation for *IRF8* expression, suggesting a bias towards myeloid differentiation is already initiated within the first 24h of IFNγ stimulation (Figure 5I).

### IFNγ-induced, differentiation-primed AML cells facilitate disease progression in the presence of anti-tumor T cell responses

Injection of unfractionated, IFNγ or PBS pre-treated NP AML cells in immunodeficient NSG mice demonstrated that IFNγ-exposed AML cells had reduced, but not completely lost, leukemia propagating capacity (Figure 6A and 6B). This was consistent with the *in vitro* data (Figure 5B-D) and suggested that the stem/progenitor AML cell compartment was reduced but not completely eliminated following exposure to this cytokine. Disease infiltration positively correlated with the injected dose of NP AML cells in both conditions.

**Figure 6.**
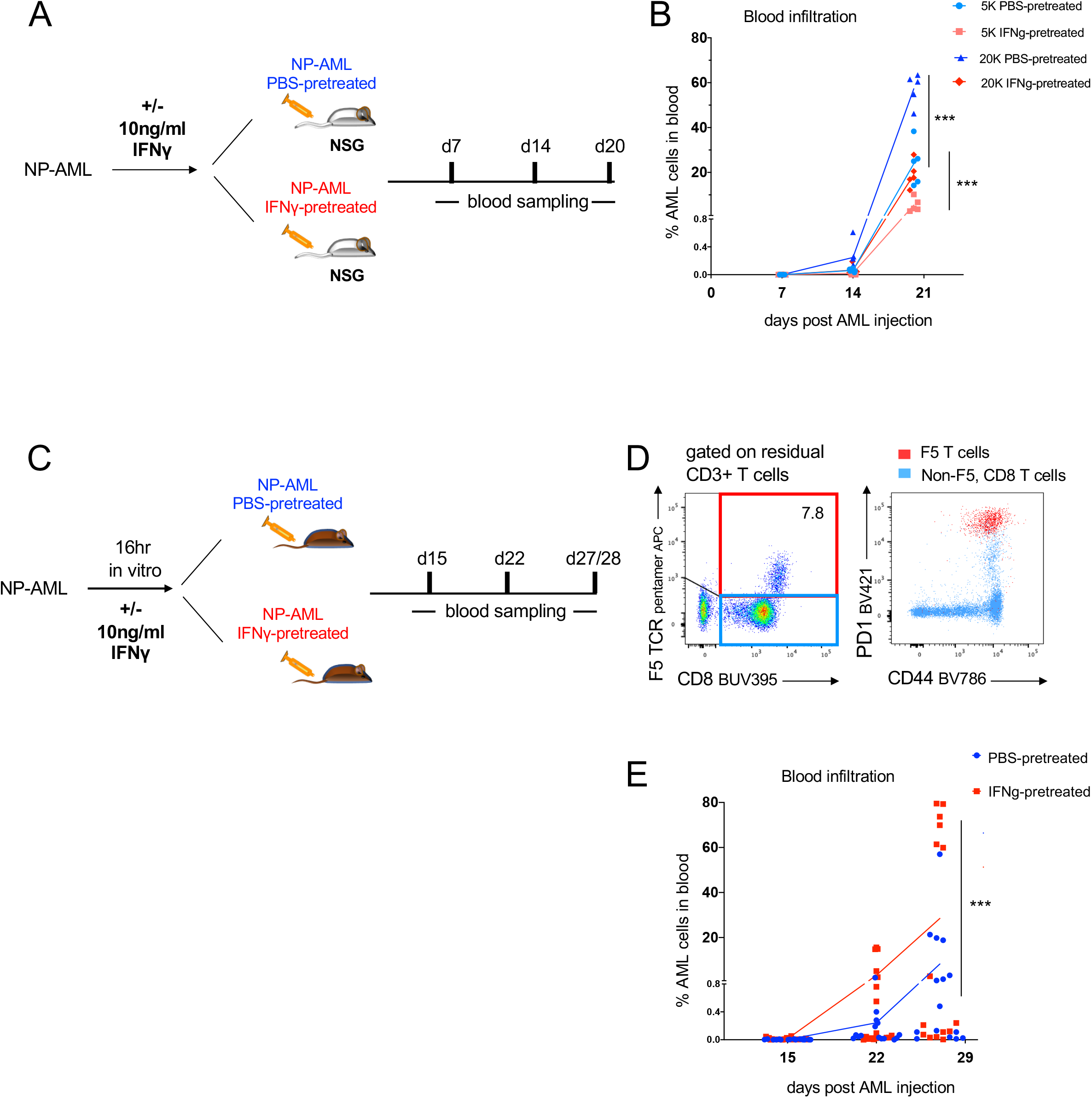
IFNγ-induced PDL1^hi^ AML cells facilitate disease progression in the presence of anti-tumor T cells responses. (A) Schematic diagram of experimental setup. NP-AML cells were pre-treated with PBS or 10ng/ml IFNγ for 48hrs and injected into NSG mice (5000 or 20 000 cells per mouse), to assess their leukemia propagating capacity. Blood was sampled for disease progression at 7, 14 and 20 days post-AML injection. n= 5 mice per group. (B) Graph summarising % leukemia infiltration in the blood of NSG mice injected with PBS or IFNγ pre-treated AML cells as described in (A). Each dot represents individual mice data, lines connect mean infiltration at each timepoint, in each group. T-tests at each time point used for statistical significance. (C) Schematic diagram of experimental setup. NP-AML cells were pre-treated with PBS or 10ng/ml IFNγ in culture overnight and injected in immunocompetent C57Bl/6 mice (100 000 cells/mouse). Blood was sampled at 15, 22, 27 days post-AML injection, to monitor disease progression. PBS pre-treated AML cells, n=15 recipient mice. IFNγ pre-treated AML cells, n= 17 recipient mice. (D) Representative FACS plot of blood samples from fully infiltrated leukemic mice from (C) stained with F5-pentamer, and antibodies against CD3, CD8, CD44 and PD1. F5-specific T cells (red gate) were PD1 and CD44 double positive. (E) Percentage blood infiltration of mice receiving PBS pre-treated or IFNγ pre-treated AML cells, at 15, 22, and 27 days post-AML injection. Each dot represents an individual mouse. Lines connect mean infiltration at each time point, in each group. T tests at each time point, with Holm-Sidak correction, used for statistical significance.

Finally, we tested whether the shift towards AML differentiation following IFNγ exposure could paradoxically facilitate disease progression in the presence of an anti-tumor T cell response by providing a mechanism of immunoevasion. To do this, 100 000 unfractionated NP-AML cells pre-treated with PBS or IFNγ *in vitro* for 16 hrs, were injected in immunocompetent C57Bl/6 mice (Figure 6C). In fully infiltrated animals, FACS staining with an F5 pentamer detected a sizeable population of F5 T cells that all expressed high levels of CD44 and PD1.

This confirmed that rare, endogenous NP-specific T cells, which were undetectable by pentamer staining in control C57Bl/6 mice, were activated and expanded *in vivo* following an antigen-specific response against the injected NP-AML (Figure 6D). Even though this response was unable to eliminate all NP-AML cells and control leukemia growth in most mice, in its presence, IFNγ pre-treated NP-AML grew faster than PBS pre-treated NP-AML did (Figure 6E). These data highlighted that AML growth in the presence of an anti-tumor T cell response *in vivo*, does not solely depend on the intrinsic growth properties of the leukemic cells but also on their immunoevading properties. Importantly, growth and immunoevasion functions can be decoupled at the single-cell level, however these hierarchically heterogeneous leukaemic cells appear to synergize to propagate disease.

## Discussion

In this study we examined the hypothesis that in AML - a malignancy composed of hierarchically heterogeneous cells - not only LSCs, but also their progeny play a role in supporting disease propagation. We demonstrate that while proliferative, progenitor-like AML cells drive disease growth, IFNγ produced by leukemia-specific cytotoxic T cells induces their differentiation into non-cycling progeny expressing high levels of PDL1. PDL1^hi^ AML cells are unable to propagate leukemia themselves, however, promote disease progression in the presence of anti-tumor T cell responses. Our work suggests a possible important role for the heterogeneous LSC progeny, with different AML cells exhibiting distinct functions and synergizing with LSCs to establish disease despite immune-driven stress.

Both inter-patient heterogeneity in the immune landscape of AML and correlations between PDL1 expression and worse clinical outcomes are becoming increasingly obvious (Antohe et al., 2020; Chen et al., 2020; Vadakekolathu et al., 2020; Williams et al., 2019). Here, we document dynamic changes in the spatial distribution of the BM T cell compartment in leukemic mice, as the disease progresses. The appearance of BM T cell clusters, either in direct contact with AML foci or in areas free of leukemia, at early/intermediate disease infiltration and the high proportion of cytotoxic, CD8 BM T cells expressing PD1 and CD44 in heavily infiltrated animals suggest that anti-tumor T cell responses take place, and these appear to initially control leukemia growth in immunocompetent mice. At early/intermediate disease we observed a transient increase of PDL1 expression levels on AML cells. Consistent with our data, a recent study combining targeted immune gene expression with multiplexed spatial profiling of BM from AML patients revealed immune TME heterogeneity and showed higher levels of PDL1 expression in immune-infiltrated BM areas exhibiting higher numbers of cytotoxic CD8 T cells and an IFNγ signature, than in CD3-deplete areas (Vadakekolathu et al., 2020). As these findings provide evidence of T cell responses shaping the disease locally, it is interesting to note that in our flow cytometry-based analyses we primarily observe the increase in PDL1 expression among AML cells in blood. Even with recent evidence that IFNγ secreted by T cells can diffuse up to 800μm within the TME (Hoekstra et al., 2020; Thibaut et al., 2020), the localized T cell responses will only impact small portions of the leukemic BM to increase PDL1 expression. It is likely that differentiation towards the patrolling monocyte fate (Bianchini et al., 2019) and low CXCR4 expression predispose PDL1^hi^ AML cells to egress into the circulation, however this happens after they immunosuppress anti-tumor T cells within the leukemic BM.

In our study, IFNγ-induced PDL1 upregulation, known to provide an immunosuppressive advantage in a variety of tumors (Dong et al., 2002; Freeman et al., 2000; Taube et al., 2012), is coupled with AML cell differentiation, reduced proliferation, and reduced leukemia-propagating capacity. These changes are in line with the cytostatic/ anti-proliferative effects of IFNγ on some cancer cells (Castro et al., 2018), and may explain the enrichment in PDL1^hi^ AML cells in both blood and bone marrow following *in vivo* chemotherapy treatment. Differentiated PDL1^hi^ AML cells likely have a limited lifespan, raising the question whether they would be clinically important. Our data supports this to be the case, as IFNγ pre-treated NP-AML proved to have a growth advantage over PBS pre-treated NP-AML, upon transplantation into immunocompetent mice that are able to mount an anti-tumor response against the NP-model antigen.

It is becoming increasingly obvious that T cell function following chemotherapy impacts on clinical outcome (Knaus et al., 2018). At the same time, chemotherapy in AML serves as a conditioning regime for both T cell immunotherapies and allogeneic hematopoietic stem cell transplantation (Dohner et al., 2017), with the curative effect of the latter also relying on the T cell driven graft-vs-leukemia response. Thus, we propose that the composition and immunomodulatory functions of chemoresistant AML cells are likely to impact the relapse risk. While the development of new AML treatments rightly focuses on the effective targeting of chemoresistant LSCs, and differentiation is generally a welcomed outcome, our study alerts against overlooking the possible impact of differentiated immunosuppressive AML cells.

More broadly, our work demonstrates that hierarchical heterogeneity in cancer can contribute to disease progression. The diverse immunomodulatory properties of hierarchically heterogeneous malignant cells might be one way by which heterogeneity facilitates tumor growth. In colon cancer and oligodendrogliomas, developmental hierarchies are being confirmed with the advancement of single cell transcriptomics (Dalerba et al., 2011; Tirosh et al., 2016). While the presence of hierarchical heterogeneity is recognized, the functional involvement of the differentiated populations, in terms of their interactions with the immune, vascular and stromal components of the tumor microenvironment, and their contribution to disease progression is understudied. Should these interactions prove to be important, it will be critical to investigate if the generation of differentiated cancer progeny in other cancers is dynamically regulated by microenvironmental changes similar to in AML.

---

**Figure S1.**
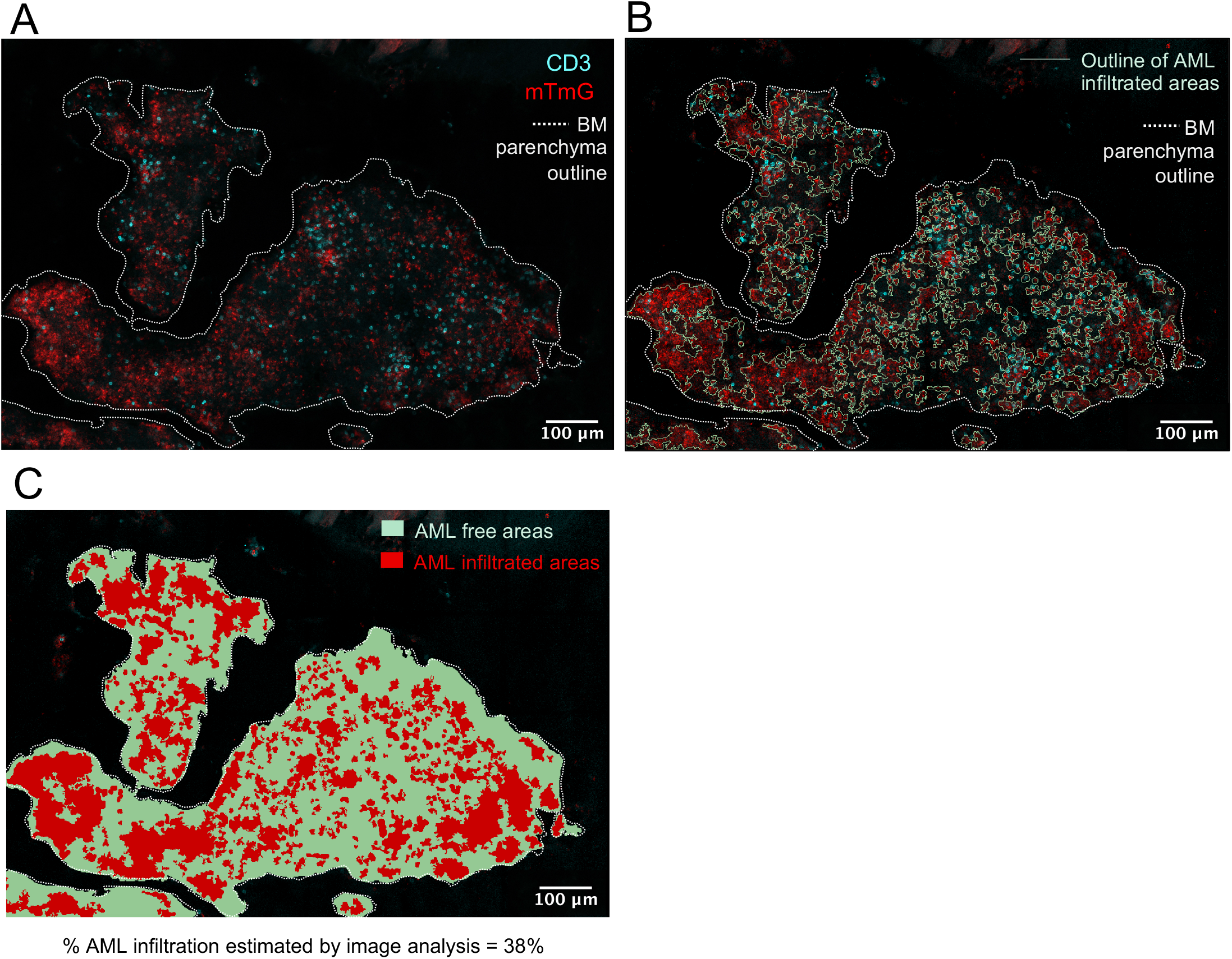
Percentage BM AML infiltration estimation by image analysis. (A) Maximum intensity projection of a tile scan image of a representative 20μm-thick section of leukemic mouse sternum, stained with anti-CD3 antibody (cyan). Leukemic cells express tomato FP (red). The dotted line indicates the edges of BM tissue; bone is not shown. (B) Same maximum intensity projection as in (A) with the tissue boundaries and AML infiltrated areas outlined in QuPath. (C) Outlined AML infiltrated areas and AML-free areas filled in red and green colours, respectively, and percentage red area (% AML infiltrated area) calculated in QuPath. In example shown, % AML infiltration estimated by image analysis was 38% and % AML infiltration by FACS in long bones (femurs, tibias, pelvic bones) was 44%.

**Figure S2.**
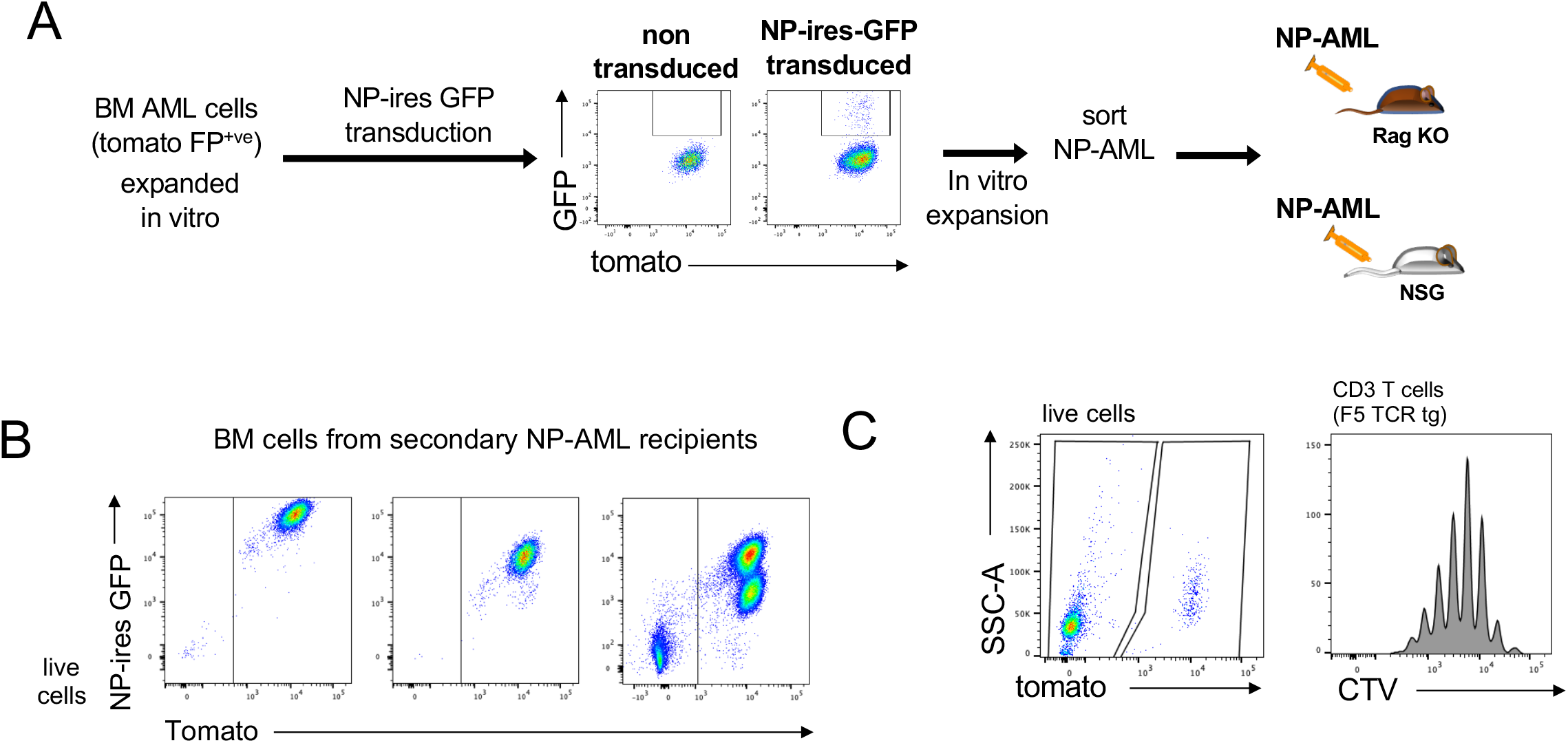
Generation of NP-expressing AML model. (A) Primary AML cells were transduced with retroviral particles encoding the NP-model antigen and GFP. NP-transduced cells were FACS purified based on high levels of GFP expression and were expanded *in vivo* in secondary Rag1 KO or NSG recipients. (B) Representative FACS plots of BM cells from three NP-AML recipient animals. NP-ires-GFP expression on AML cells varied between individual animals, however either all or a significant proportion of cells expressed higher GFP levels than untransduced ones. (C) Representative FACS plots of CTV-labelled F5 TCRtg splenocytes (tomato-) in culture with NP-AML (tomato+), plated at 2:1 ratio and cultured for 6 days. The FACS plot on the left shows all live cells and the histogram on the right shows CTV dilution among tomato^-ve^ F5 T cells (gated on tomato^-ve^, CD3^+ve^ cells), indicating T cell proliferation.

**Figure S3.**
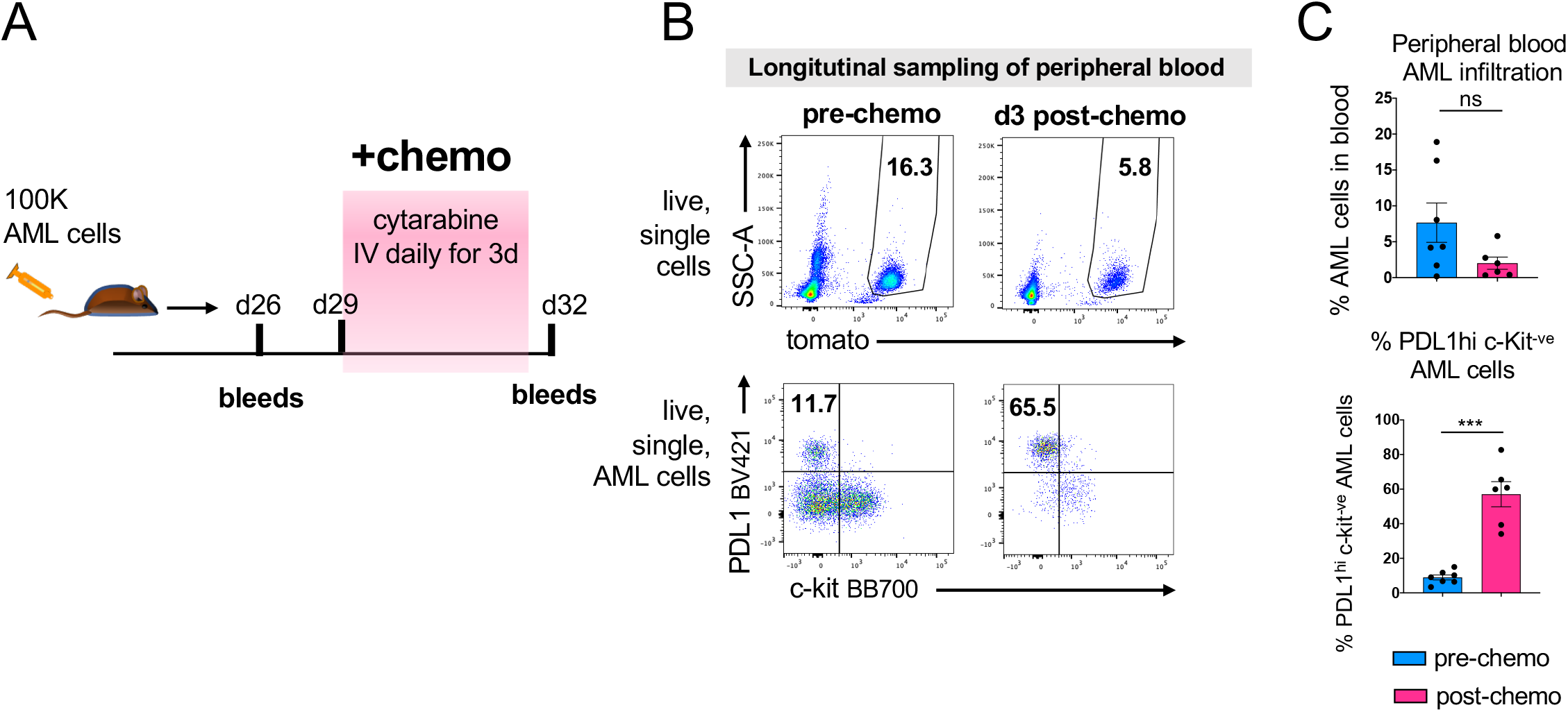
Chemoresistant AML is highly enriched in PDL1hi AML cells. (A) Schematic diagram of experimental setup. 100 000 AML cells were injected in C57Bl/6 recipients. Blood was sampled on d26 post-AML injection, mice were then administered with IV cytarabine for 3 days on d29, d30 and d31 and blood was sampled again on d32 to monitor AML infiltration and phenotype AML cells for PDL1 and c-Kit expression. n= 7 C57Bl/6 leukemic mice (1 out of 7 mice had to be sacrificed prior to the d32 timepoint, due to poor health) (B) Representative FACS plots of blood collected from leukemic mice. Top row shows % infiltration on d26 and d32 (as described in A). Bottom row shows PDL1 and c-Kit phenotype of AML cells in the same subject at these time points. (C) Top graph summarises the percentage infiltration in the blood of leukemic mice on d26 and d32 (as described in A). Bottom graph summarising the percentage of PDL1^hi^ c-Kit^-ve^ AML cells on d26 and d32 (as described in A). Each dot represents data from individual mice. Paired t-test used for statistical significance, error bars: means ± SEM.

**Figure S4.**
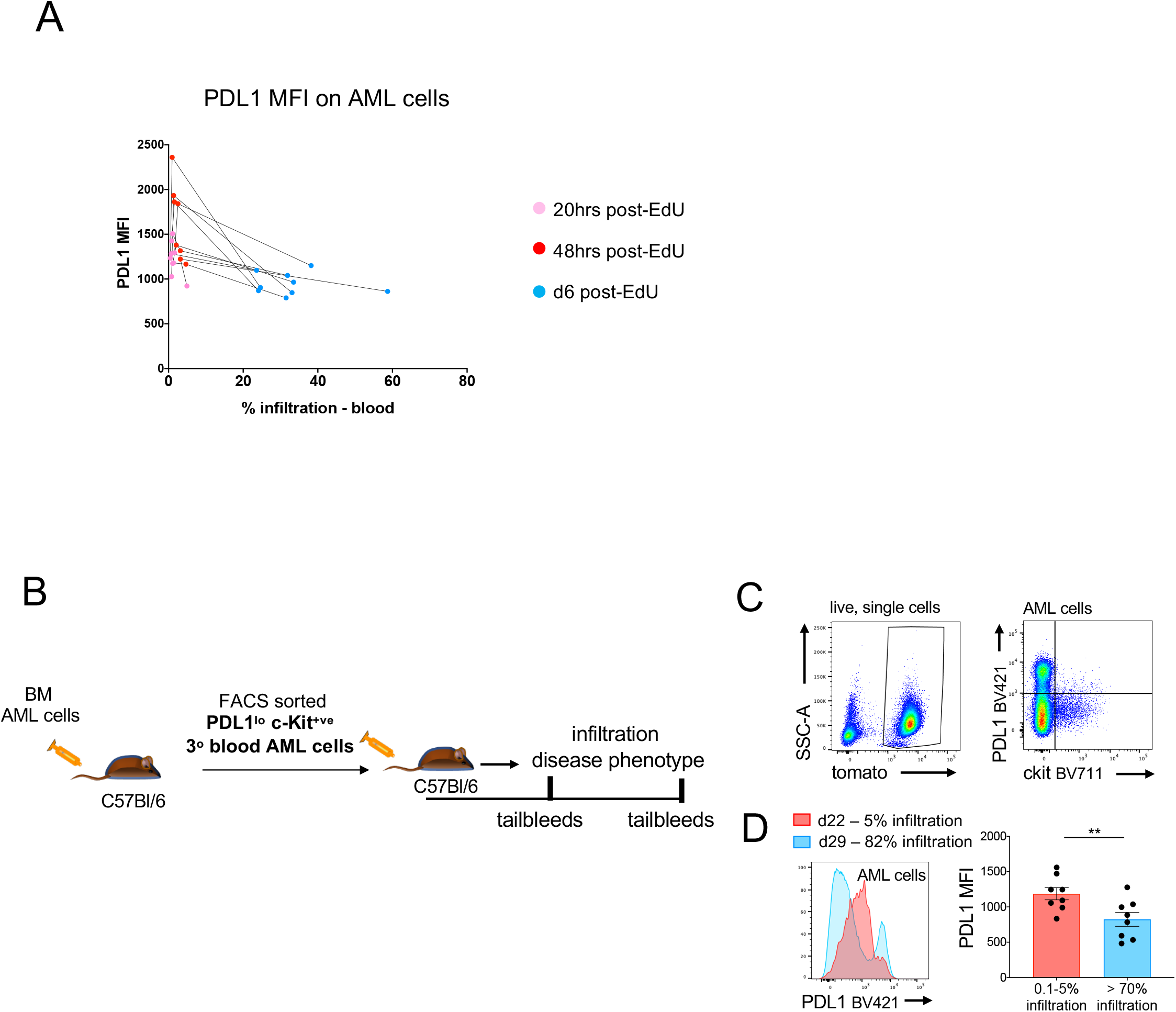
Transient increase in PDL1 expression levels among AML cells, in lineage tracing experiments. (A) Graph showing PDL1 MFI values on blood AML cells for each mouse at 20hr, 48hr and d6 post-EdU pulse, in the EdU pulse-chase experiment in relation to % AML infiltration in the blood. Each dot represents data from individual mice and data from each mouse are connected with a line. (B) Schematic diagram of experimental setup. Primary BM AML cells were serially transplanted into C57Bl/6 secondary and subsequently tertiary recipients. PDL1^lo^ c-Kit^+ve^ AML cells were FACS sorted from the pooled blood of tertiary recipients and transplanted into quaternary C57Bl/6 recipients. Blood was sampled to monitor disease growth and AML cell phenotype in quaternary recipients. (C) Representative FACS plots of blood samples from quaternary C57Bl/6 AML recipients, as described in (A), showing PDL1 and c-Kit expression (right-side plot) on AML cells. (D) Histogram overlay of PDL1 expression on blood AML cells in early and late infiltration and graph showing PDL1 MFI at early-intermediate (0.1<AML blood infiltration<5) and late disease (AML blood infiltration>70%) infiltration. Graph summarises, PDL1 MFI on AML cells from individual mice at AML infiltration<5% and then in the same mice when AML infiltration >70%. Paired t-tests used for statistical significance, error bars: mean± SEM.

## Materials and Methods

### Mice

Animal work was performed in accordance with the animal ethics committee (AWERB) at Imperial College London and the Francis Crick Institute, UK and following UK Home Office regulations (ASPA, 1986). NOD-scid IL2Rgamma^null^ (NSG), F5 Rag1KO CD45.1 (C57Bl/6) transgenic (F5 TCRtg), OTI Rag1KO (C57Bl/6) transgenic (OTI TCRtg), Rag1KO (C57Bl/6) (RAG KO) and C57Bl/6 mice were bred and housed at the Francis Crick Institute, in individually ventilated cages. Aged-matched, female mice> 6 weeks old were used for all experiments.

### AML experimental model

Primary AML cells were generated from granulomyelocytic progenitors from mT/mG or PU.1-YFP mice (C57Bl/6 background) using MLL-AF9-IRES-GFP retroviral particles as described (Duarte et al., 2018). BM cells and splenocytes were harvested from primary mice, when fully infiltrated with leukemia (BM AML infiltration>90%), and were cryopreserved as individual batches of primary AML cells (Duarte et al., 2018). Primary AML cells were thawed, washed and resuspended in PBS and up to 100 000 cells were injected into secondary syngeneic C57Bl/6 recipients or immunodeficient Rag KO or NSG mice, as described in individual experiments.

To generate AML cells expressing the NP-model antigen, PlatE adherent packaging cells were transfected using Calcium phosphate precipitation (Invitrogen) with the pMP71 NP-ires-GFP construct (kind gift from H.J.Stauss). Viral supernatant was harvested from the transfected cells and used to transduce primary BM AML cells by spinfection on retronectin-coated (Takara-Bio Inc) plates. The transduced NP-AML cells were expanded *in vitro* and GFP bright cells, which expressed the model antigen, were purified by FACS and injected intravenously for *in vivo* expansion in Rag KO and NSG mice. Spleens and BM were harvested from fully infiltrated animals and single cell suspensions from each mouse were frozen down as individual batches of NP-AML cells.

### Chemotherapy/ Drug Treatments

Chemotherapy was administered when AML was detected in the blood of mice, which corresponds to BM infiltration over 30% (Akinduro et al., 2018). 100mg/kg cytarabine was administered IV for 3 days and in specified experiments in combination with 3mg/Kg doxorubicin. Both drugs were obtained from Sigma Aldrich.

### EdU and BrdU administration

EdU (1mg/mouse) and BrdU (2mg/mouse) were administered to leukemic mice via tail vein injections. In the Edu/BrdU dual pulse chase experiments, these were administered 2hr apart and 30min following the BrdU injection mice were culled, and BM, spleen and blood were harvested for immunophenotypic and cell cycle analysis of leukemic cells in these tissues.

In the EdU pulse-chase experiment, a single pulse of EdU was followed by longitudinal sampling of blood at 20hr, 48hr and 6days post-EdU administration, for immunophenotypic and Edu label-dilution analysis in blood AML cells.

### Tissue harvest and single cell suspension preparation

Blood was sampled either by venipuncture of the mouse tail, or through cardiac bleeds (following CO2 administration). To prepare single cell suspensions of splenocytes, freshly harvested spleens were mashed and passed through a 70-μm cell strainer. To prepare single cell suspensions of BM cells, femurs, tibias and pelvic bones were crushed with a pestle and mortar in RPMI medium supplemented with 10% FBS and passed through a a 70-μm cell strainer. Red blood cells were removed by isotonic lysis with ammonium chloride. Cells were resuspended in FACS buffer (PBS, 2% FCS) for counting, immunolabelling and FACS analysis.

### Murine AML cell culture

AML cells were plated at 0.25×10^6^/ml in RPMI medium, supplemented with 10% FBS, 1% penicillin/streptomycin, 1% L-glutamine and 10ng/ml recombinant murine SCF, 6ng/ml recombinant murine IL-3 and 10ng/ml recombinant human IL-6. When specified, 10ng/ml recombinant murine IFNγ was added to the cultures. All cytokines were purchased from Peprotech. For co-culture experiments, 100 000 NP-AML AML cells were plated in the above media, together with 200 000 splenocytes from F5 or OTI TCR transgenic mice, in 96-well u-bottom plates, in 200μl/well. In some experiments, AML cells were cultured in the above media with the addition of conditioned media (1:1 volume) from F5:NP-AML cocultures, or OTI:NP-AML cocultures, in the presence or absence of an IFN γ blocking antibody (R4-6A2 clone, NA/LE, BD Biosciences). Cells were incubated at 37°C, 5% CO2.

### Chemotherapy treatment of Cell Trace labelled AML cells

BM, spleen and blood cells were harvested from leukemic mice and labelled with 1μM Cell Trace Violet, or 0.2μM Cell Trace Red (Thermofisher Scientific), as per manufacturers protocol. The cells were then plated at 0.25×10^6^/ml in the presence or absence of 3μM cytarabine and 450nM doxorubicin and incubated at 37°C, 5% CO2. FACS analysis for immunophenotype and cell trace dilution were performed at the indicated time points.

### Colony forming Assays

AML cells were mixed in 1ml of MethocultTM (M3434; StemCell Technologies) and plated in 6-well plates (1ml/well). In some experiments, AML cells were FACS sorted based on their immunophenotype and directly into 6-well plates containing 1ml of MethocultTM per well. Cultures were incubated at 37°C and quantified on day6 to day9. In specified experiments, 10ng/ml of IFNγ or PBS was added to the cultures.

### Primary AML samples and culture

AML samples were obtained with informed consent at St Bartholomew’s Hospital (London, UK). The cells were collected and frozen at diagnosis. Details of patient samples are provided in Supplementary Table 1. AML mononuclear cells were isolated by centrifugation using Ficoll-Paque (GE Healthcare Life Sciences). Immuno-magnetic T and B cell depletion was performed using the EasySep magnetic system (StemCell Technologies), prior to setting up *in vitro* cultures. Cells were plated at 200,000 cells/ml Stem Span medium (StemCell Technologies) supplemented with 3GT cytokines (20ng/ml IL-3, 20ng/ml G-CSF, 20ng/ml TPO overnight, prior to the addition of 10ng/ml IFNγ or PBS control. All cytokines were from Peprotech. Cells were incubated at 37°C, 5% CO2. 24hr after the addition of IFNγ or PBS, cells were harvested for qPCRs. 3days after the addition of IFNγ or PBS, cells were harvested for FACS analysis.

### Flow Cytometry and cell sorting

For the immunophenotypic analysis of T cells and AML cells, the following fluorochrome-conjugated primary antibodies specific to mouse were used: CD3e (145-2C11 or 17A2), CD4 (GK1.5), CD8a (53-6.7), CD44 (IM7), PD1 (29F.1A12 or J43), PDL1 (M1H5 or 10F.9G2), c-Kit (2B8), CD11b (M1/70), CD11c (HL3 or N418), Ly6G (1A8), Ly6C (HK1.4), MHC II (M5/114.15.2), CX3CR1 (SA011F11), CXCR4 (L276F12), Ki67 (11F6 or B56), BrdU (MoBu-1). Antibodies were purchased from BDBioscience, Biolegend and eBioscience. Live and dead cells were distinguished using 7AAD (Biolegend or BDBiosciences), or fixable viability dyes 780, or BV510 (BD Biosciences).

Calibrite beads (BD Biosciences) were used to determine absolute cell counts. Cells were analyzed with a BD Symphony or LSR Fortessa, and sort purified using a FACSAria (BD Biosciences) and data were analyzed with FlowJo (TreeStar).

### Immunofluorescence of non-decalcified bone sections

Tibias and sterna from healthy and leukemic mice were harvested and fixed overnight in 4% PFA, at 4°C. The bones were then washed in PBS and taken through a sucrose gradient (10%, followed by 20%, and finally 30% sucrose in PBS) over 48-72hrs. They were then frozen in optimal cutting temperature (OCT) compound (TissueTek) and kept at −80°C. 20μm sections were obtained using a cryostat (Leica) and the Cryojane tape transfer system (Leica microsystems). Slides were stored at −80°C. For staining, slides were thawed, rehydrated in PBS, and permeabilized in 0.1% Triton X-100. They were blocked in 5% goat serum and incubated with CD3e BV421 or AF647 antibody (clone 17A2, 1:100) in 0.1% Triton X100, supplemented with 5% goat serum, for 2hr at RT, in the dark. Slides were then washed in 0.1% Triton X-100 and mounted with Prolong Diamond antifade (Invitrogen). Stack images were obtained with a z-step of 2μm using a Zeiss LSM880 invert confocal microscope and analyzed using Fiji/ImageJ and QuPath.

### Image processing and quantification

Zen black (Zeiss, Germany) software was used to acquire and stitch three-dimensional overlapping tiles to form composite BM tilescans. These were visualized and processed into two-dimensional maximum projection images in Fiji software (ImageJ 1.50). Images were then uploaded into QuPath software (QuPath version 0. 2. 1) (Bankhead et al., 2017). The total area of the BM tissue was calculated by manually outlining the ROI with the built-in *wand tool*. AML infiltration was estimated as a % measure of mTomato^+^ or YFP^+^ cells per overall tissue area, using the built-in *wand tool* in QuPath software to highlight mTomato^+^ or YFP^+^ cells (Figure S1). To count T cell clusters within BM tissue, maximum projection images were divided into 0.25mm^2^ regions covering the BM parenchyma and clusters were counted per area. A T cell cluster was defined as >20 cells/2000μm^2^, and clusters were identified using a custom-made Fiji Macro (available on request) and results were validated manually. T cell size was measured in QuPath following manual segmentation. Total numbers of T cells were manually counted using the built-in *points* cell counting tool. Raw data was processed in Microsoft Excel.

### Statistical analysis

Statistical analysis was performed using GraphPad Prism (GraphPad Software).

Group means were compared using the unpaired Student’s t-test, with Holm-Sidac correction for multiple comparisons. When samples compared where paired, individual values were compared using the paired Student’s t-test. For all data, differences were considered significant whenever p<0.05. * p <0.05, ** p< 0.01, *** p < 0.001, **** p<0.0001. Number of animals and samples used specified in Figure legends.

## Author contributions

CP, CLC, WG and DB were responsible for the concept and design of the study. CP, WG, SGA and SG performed experiments, data analysis and interpreted the data. CG, FB, TW, RK, MH, NMSP, KS, HHE, JH, CC, DS, FHY, performed experiments and data analysis. GS developed the image analysis methodology and performed image analysis. WG, HJS, RC and DB provided expertise and critical feedback. All authors contributed to the manuscript. CP and CLC wrote the manuscript. CP, CLC and DB acquired the funding supporting this work.

## Acknowledgements

We thank Tiago Cunha Luis and Caetano Reis e Sousa for critical discussions and feedback, the Francis Crick Institute FACS, Imaging, Biological Research Facilities, and the Imperial College FILM facility for technical support. We thank Stephen Rothery (Imperial College FILM facility) for writing the Fiji Macro to automatically identify T cell clusters. The manuscript was edited by Life Science Editors.

---

**Table S1.** Details on patient samples, treated with IFNγ in vitro

|  | Genetics | FAB classification |
| --- | --- | --- |
| Patient 1 | ADD(21)(p11.2) | M5a |
| Patient 2 | MLL-AF6 | M4 |
| Patient 3 | MLL-AF9 | M5a |
| Patient 4 | MLL-AF9 | M5 |

**Table S2.**
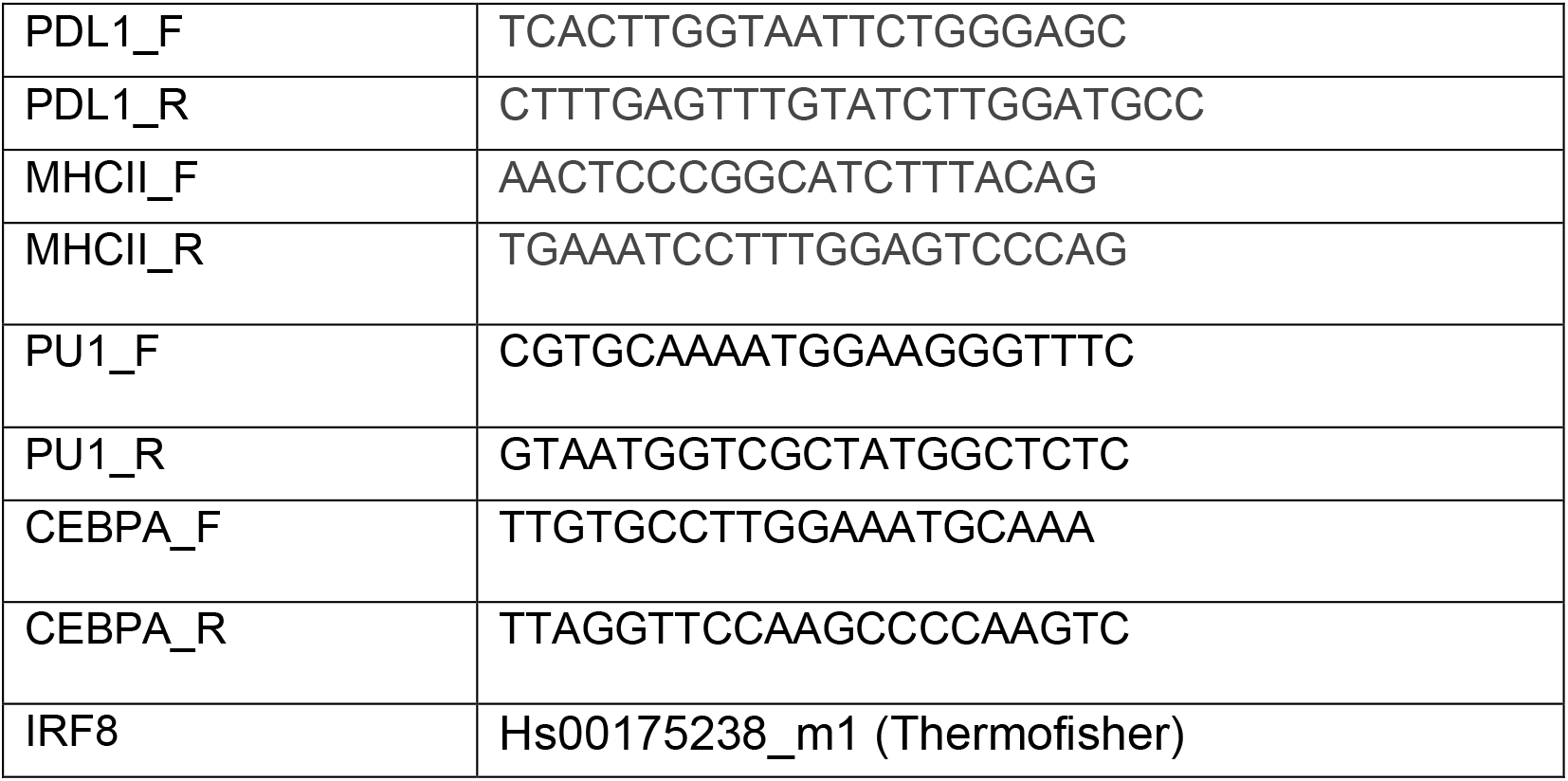
List of qPCR primers

